# Converging pathways: shared brain circuitry engaged by monoaminergic antidepressants, ketamine and psilocybin

**DOI:** 10.1101/2025.05.26.655791

**Authors:** Kadeem Joseph, Janelle Collins, Thomas Genovese, Myran Maxwell, Jeffrey A. Lieberman, Pavel Osten

## Abstract

Ketamine has transformed depression treatment by providing therapeutic relief within a single day, unlike monoaminergic antidepressants that require weeks to take effect. Here, we conducted whole-brain screening in mice to compare drug-evoked c-fos expression—acting as a marker of brain activity leading to protein synthesis-dependent forms of plasticity—following treatment with monoaminergic antidepressants, ketamine and psilocybin. Our findings reveal a shared limbic brain circuit comprising subcortical and frontal cortical regions, with a key distinction: c-fos-based activity in the prelimbic and infralimbic frontal cortex—areas strongly implicated in depression—was acutely induced by ketamine and high-dose psilocybin, but emerged only after chronic dosing with the selective serotonin reuptake inhibitor fluoxetine or psilocybin microdosing. These results suggest the existence of a core limbic subcortico-cortical circuit underlying antidepressant efficacy, provide mechanistic insight into the delayed therapeutic effects of monoaminergic antidepressants, and reveal a close similarity in brain activity evoked by monoaminergic antidepressants and psilocybin microdosing.

## Introduction

The monoaminergic hypothesis of depression posits that decreased serotonin, norepinephrine, and dopamine activity underlies depressive symptoms (Delgado, 2000). This hypothesis led to the development of traditional antidepressants that increase monoamine levels, including selective serotonin reuptake inhibitors (SSRIs) and serotonin-noradrenaline reuptake inhibitors (SNRIs). However, these drugs present a paradox: although they rapidly increase monoamine levels, their therapeutic effects emerge only after several weeks of treatment. This delayed onset is thought to result from gradual neuroplastic changes induced by the antidepressant treatment, such as synaptic plasticity and remodeling (Clark et al., 2009; Roiser et al., 2012).

Supporting this view, a recent positron emission tomography (PET) study in humans demonstrated a positive correlation between the duration of escitalopram (an SSRI) treatment and increased synaptic vesicle glycoprotein 2A (SV2A) density, suggesting that synaptic changes may indeed underlie the time-dependent delay in antidepressant efficacy (Johansen et al., 2023). However, the specific brain regions and circuits engaged in antidepressant-linked brain plasticity remain unknown.

In contrast, subanesthetic ketamine—a noncompetitive NMDA receptor antagonist— induces rapid and robust antidepressant effects, achieving a single-dose maximal therapeutic benefit within 24–72 hours of a single dose (Berman et al., 2000; Zarate Jr et al., 2006).

Similarly, clinical trials with psilocybin, a naturally occurring prodrug of the hallucinogenic serotonergic agonist psilocin, suggest that a single dose, typically combined with psychological support, can yield rapid and sustained antidepressant effects lasting weeks (Davis et al., 2021; Goodwin et al., 2022; Raison et al., 2023). The rapid efficacy of ketamine and psilocybin suggests mechanisms distinct from those of monoaminergic antidepressants, possibly involving stronger modulation of neural activity or targeting additional brain regions and circuits (Duman et al., 2016).

Neuroimaging studies in patients with major depressive disorder (MDD) consistently implicate abnormal activity in the medial prefrontal cortex, particularly the perigenual (pgACC) and subgenual (sgACC) anterior cingulate cortex, as central to the pathophysiology of depression (Alexander et al., 2021; Drevets et al., 2008). Subanesthetic ketamine acutely modulates activity in these regions, supporting their key role in its rapid antidepressant effects (Alexander et al., 2021; Deakin et al., 2008; Morris et al., 2020). Moreover, recent research highlights a connection between the human ACC and nucleus accumbens (NAc), revealing an expanded salience network comprising these structures in depression (Lynch et al., 2024). Together, these findings highlight the pgACC and sgACC as critical hubs in depression pathology and key targets for rapid-acting antidepressants.

To systematically investigate how different antidepressants modulate brain activity, we employed whole-brain mapping of the immediate early gene (IEG) c-fos in mice (Kim et al., 2015; Renier et al., 2016). C-fos encodes an inducible transcription factor (part of the AP-1 complex) that converts extracellular signals into new protein synthesis involved in various forms of cellular plasticity (Curran and Morgan, 1987). In this context, drug-evoked increases in c-fos-positive (c-fos+) cell counts—here referred to as “c-fos-based activity”—identify brain regions where drug-evoked activity has triggered protein synthesis associated with synaptic plasticity, remodeling, and other cellular plasticity-related processes.

We tested a range of antidepressants with established efficacy but differing mechanisms and temporal profiles: SSRIs fluoxetine and fluvoxamine, tricyclic antidepressants (TCAs) desipramine and amitriptyline, and ketamine. We also evaluated psilocybin’s potential mechanism of action in depression, including an acute hallucinogenic dose and repeated microdose. All drugs were administered at therapeutically relevant doses, based on human-equivalent dose (HED) conversions (Nair and Jacob, 2016).

Our findings identify a shared set of frontal cortical and subcortical limbic brain areas engaged by these treatments, but with distinct temporal and dose-dependent profiles across drug classes. Notably, a single treatment with ketamine or a high (hallucinogenic) dose of psilocybin acutely induced c-fos-based activity in the mouse PL and ILA cortex—homologous to the human pgACC and sgACC, which are strongly implicated in depression (Alexander et al., 2019; Alexander et al., 2021; Pizzagalli and Roberts, 2021). In contrast, the SSRI fluoxetine and a low (microdose) of psilocybin required prolonged administration to recruit these frontal cortical regions. Additionally, ketamine and high-dose psilocybin each evoked unique activity patterns in brain regions linked to their distinctive subjective effects—dissociative and hallucinogenic, respectively. These results suggest that despite differing primary pharmacological targets, diverse antidepressant therapies converge on key limbic brain circuitry to achieve antidepressant efficacy. Furthermore, parallels in brain-activation patterns between monoaminergic antidepressants and psilocybin microdosing suggest that psychedelic microdosing may exert their effects via mechanisms largely shared with conventional antidepressants.

## Methods

### Animals

Male C57BL/6 mice (8–10 weeks of age) were obtained from the Jackson Laboratory and housed in pairs at room temperature of 23 ± 2 °C, relative humidity of 60 ± 10% and artificial lighting between 7:00 and 19:00. Food and water were available ad libitum. The study will be carried out in accordance with Guidelines for Animal Care and Use at Broad Hollow Bioscience Park (BHBP). The BHBP animal facility is fully accredited by the American Association for Accreditation of Laboratory Animal Care. Animals are maintained in accordance with the applicable portions of the Animal Welfare act and the Department of Health and Human Services Guide for the Care and Use of Laboratory Animals. Veterinary care is under the direction of a consulting veterinarian boarded by the American College of Laboratory Animal Medicine. Additional veterinary staff and veterinary technicians provide a complete comprehensive program of diagnostics, preventive and clinical medicine at our facility. The consulting veterinary has authority to terminate all animal research which does not comply with current government regulations. *Drugs:* fluoxetine, fluvoxamine, desipramine, amitriptyline (all Sigma Aldrich) were dissolved in 0.3% Tween Saline, ketamine (Pfizer), psilocybin and 2C-T (Cayman Chemical) were dissolved in 0.9% Saline.

### Experimental Procedures

Adult male C57BL/6 mice were isolated (paired animals per cage with environmental enrichment to avoid stress) for 3 days prior to vehicle or test compound administration to lower the baseline variability in c-fos expression. On the experimental day between 9 to 10 am, each mouse was weighed, and vehicle or test compound was administered at selected doses by intraperitoneal (i.p.) administration. Doses were administered based on the weight of the free base. Control mice received the same volume of vehicle solution. Compound and vehicle dosing was interleaved across the test day. Mice were then immediately returned to their home cages. After a 2.5-hour period to allow for drug-evoked changes in c-fos protein expression, the mice were anesthetized with ketamine/xylazine and euthanized via transcardial perfusion with saline followed by 4% formaldehyde. The brains will then be dissected and post-fixed in 4% formaldehyde.

Brain processing, imaging and c-fos detection: C-fos expression was visualized using a modified whole-brain immunohistochemistry procedure iDISCO+ (Renier et al., 2016), with main modifications in the tissue delipidatipon step to improve antibody penetration. The brains were incubated in PBS / 0.2% TritonX-100 / 20% DMSO / 0.3M glycine at 37°C for 36h, then blocked in PBS / 0.2% TritonX-100 / 10% DMSO /6% Donkey Serum at 37°C for 2 days. Next, brains were incubated in primary anti-c-fos antibody (9F6, Cell Signaling Technology) in PBS-Tween 0.2% with Heparin 10µg/mL (PTwH) / 5% DMSO / 3% Donkey Serum at 37°C rotating for 7 days. Next, brains were washed in PTwH for 24h (5 changes of the PTwH solution over that time), then incubated in secondary antibody (donkey anti-rabbit-Alexa647 at 1/500th in PTwH / 3% Donkey Serum) rotating at 37°C for 7 days. Finally, the samples were washed in PTwH for 1d before clearing and imaging.

The immunolabeled brains were cleared as follows: brains were dehydrated in 20% methanol (in ddH2O) for 1h, 40% methanol / H2O for 1h, 60% methanol / H2O for 1h, 80% methanol / H2O for 1h, and 100% Methanol for 1h twice. Brains were then incubated overnight in 1 volume of Methanol / 2 Volumes of Dichloromethane (DCM, Sigma 270997-12X100ML) until they sank at the bottom of the vial. Methanol was washed for 20min twice in 100% DCM. Finally, brains were incubated (without shaking) in DiBenzyl Ether (DBE, Sigma 108014-1KG) until clear (about 30min) and then stored in DBE at room temperature.

### Computational and Statistical Methods

Whole-brain distribution of activated c-fos+ neurons in the test compound-and vehicle-treated groups was imaged by light-sheet fluorescent microscopy (LSFM) as published (Renier et al., 2016). The brains were imaged in sagittal orientation (right lateral side up) on a light-sheet fluorescence microscope (Ultramicroscope II, LaVision Biotec) equipped with a sCMOS camera (Andor Neo) and macrolens equipped with dipping cap. The samples were scanned with a step-size of 5-micrometer using the continuous light-sheet scanning method at 640 nm for c-fos signal channel and 480-nm for anatomical registration channel. The activated c-fos+ neurons were automatically identified by custom computational algorithms and visualized in 3D (see the entire pipeline in Supplemental Figure 1). The datasets were warped in 3D by affine and B-spline transformation to an average Reference mouse brain generated from 8-week old C57BL/6 brains.

Statistical comparisons between vehicle and drug-treated groups were run based on evenly spaced voxels and anatomical atlas ROI’s Voxels are overlapping 3D spheres with 100 μm diameter each and spaced 20 μm apart from each other. The cell counts at a given location, *Y*, are assumed to follow a negative binomial distribution whose mean is linearly related to one or more experimental conditions, *X:* E*[Y]=α+βX*. For example, when testing a control group versus an experimental group, the *X* is a single column showing the categorical classification of mouse sample to group id, i.e. 0 for the control group and 1 for the experimental group (O’Hara and Kotze, 2010; Venables and Ripley, 2002). The maximum likelihood coefficients *α* and *β* was found through iterative reweighted least squares, obtaining estimates for sample standard deviations in the process, from which the significance of the *β* coefficient was obtained. A significant *β* means the group status was related to the cell count density at the specified location. The z-values correspond to this *β* coefficient normalized by its sample standard deviation, which under the null hypothesis of no group effect, has an asymptotic standard normal distribution. The p-values give the probability of obtaining a *β* coefficient as extreme as the one observed by chance assuming this null hypothesis is true. To account for multiple comparisons across all voxel/ROI locations, the p-values were thresholded by false discovery rates with the Benjamini-Hochberg procedure (Benjamini and Hochberg, 1995).

## Results

### Study design

The whole-brain c-fos mapping pipeline and computational methods used to analyze the data, previously described by us and colleagues (Azevedo et al., 2020a; Azevedo et al., 2020b; Kim et al., 2015; Renier et al., 2016), enabled unbiased screening of drug-evoked responses across the mouse brain at single-cell resolution (Methods; Supplemental Figure 1). In brief, c-fos+ cells from each mouse brain were automatically detected and co-registered to a 3D reference brain segmented into ∼7.8 million voxels (150 μm each). Drug-induced changes in c-fos+ cell counts were then calculated by voxel-wise statistical comparisons between vehicle-treated (control) and drug-treated groups, using negative binomial regression with false discovery rate (FDR) correction for multiple comparisons. The results are visualized as color-coded maps of statistically significant differences, with increases in c-fos+ cell counts shown in red and decreases in green, overlaid on the grayscale reference brain (see representative coronal sections in Figure 1–Figure 3 and Supplemental Figures 2–4). In addition, the reference brain was co-registered with a digital anatomical atlas (Allen Mouse Brain Atlas (Dong, 2008)), enabling a complementary region-of-interest analysis. Drug-induced changes in c-fos+ cell counts were quantified for each annotated brain region, also using negative binomial regression with FDR correction (see bar graphs in Figure 1–Figure 3 and Supplemental Figures 2–4).

**Figure 1.**
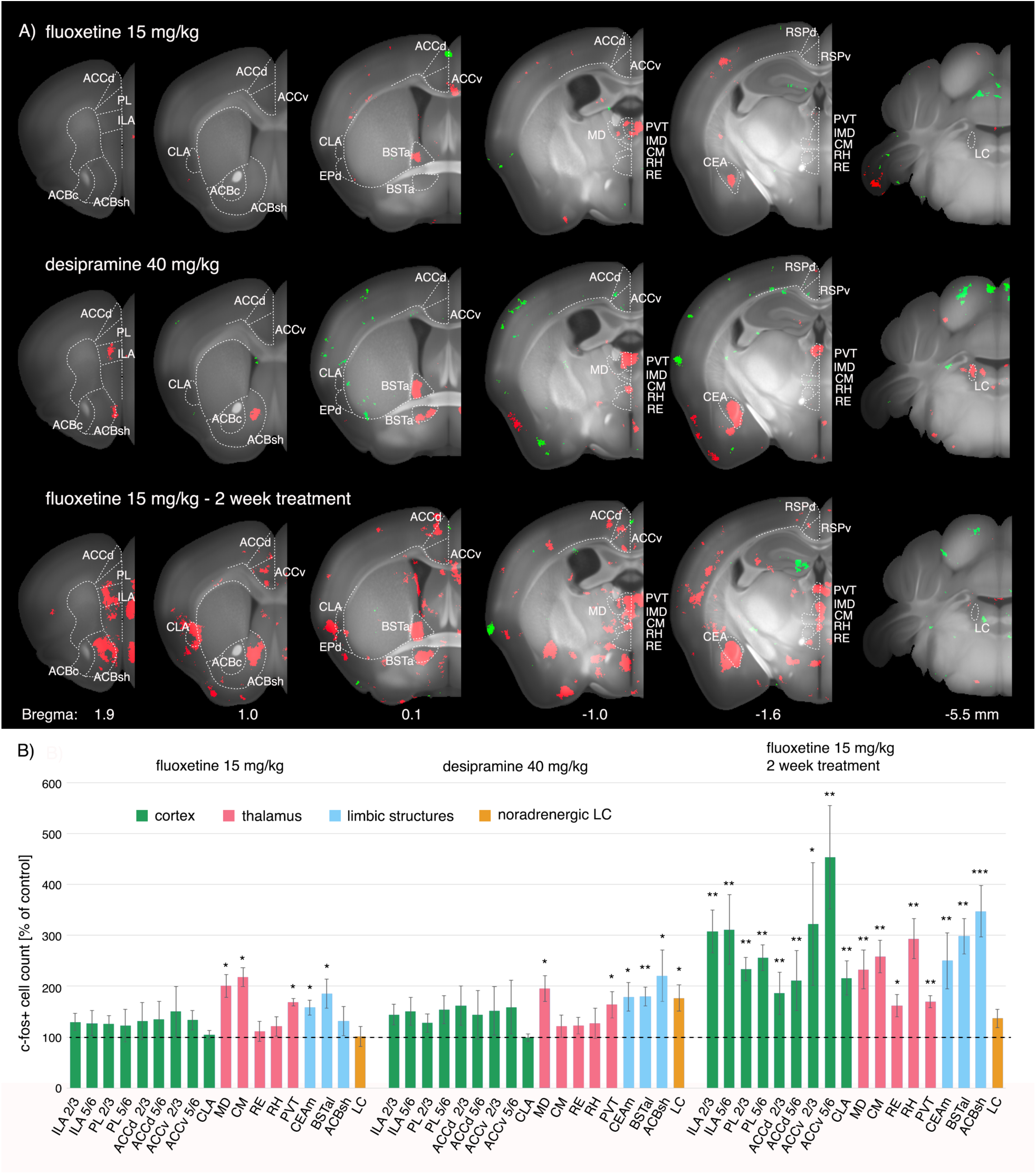
Brain activity evoked by single and chronic treatment with monoaminergic antidepressants. (**A**) Brain regions showing statistically significant increases in c-fos expression (red) are overlaid on selected coronal sections of the 3D reference mouse brain (grayscale). Green indicates areas with decreased c-fos expression; these decreases were sparse and are likely not biologically meaningful. *Top:* Activation pattern evoked by a single dose of the SSRI **fluoxetine** (15 mg/kg). *Middle:* Pattern evoked by a single dose of the TCA **desipramine** (30 mg/kg). *Bottom:* Pattern evoked by a chronic (two-week) fluoxetine treatment (15 mg/kg daily). Note the more robust subcortical activity evoked by a single dose of desipramine compared to fluoxetine, as well as the substantially increased frontocortical and thalamic activity observed after two weeks of fluoxetine relative to either acute fluoxetine or desipramine. (**B**) Quantification of the c-fos+ cell count changes corresponding to the regions shown in (A), expressed as percent change in drug-treated vs. vehicle-treated groups (mean ± SEM). Statistical significance was assessed by negative binomial regression with FDR correction for multiple comparisons: q* < 0.05, q** < 0.01, q*** < 0.001. See Methods for anatomical region abbreviations.

### Shared c-fos-based brain activity by monoaminergic antidepressants

We first mapped brain activity induced by single doses of the monoaminergic antidepressants. The initial set of drug-screening experiments compared the effects of single treatments with the SSRIs fluoxetine (15 mg/kg) and fluvoxamine (60 mg/kg), and the tricyclic antidepressants (TCAs) desipramine (40 mg/kg) and amitriptyline (15 mg/kg). Whole-brain voxel-wise analysis of drug-evoked activity revealed a discrete yet consistent subcortical activation pattern that was shared across all four antidepressants (Figure 1; Supplemental Figure 2; Methods). Specifically, each treatment induced statistically significant increases in c-fos+ cell counts in the mediodorsal thalamus (MD), paraventricular thalamus (PVT), central amygdala (CEA), and the anterior part of the bed nucleus of the stria terminalis (BSTa) (Figure 1; Supplemental Figure 2; Supplemental Table 1). Additionally, the TCA’s desipramine and amitriptyline—but not SSRI’s fluoxetine or fluvoxamine—induced hotspot-like c-fos-based activity in the medial part of the shell of the nucleus accumbens (NAc-sh) and the noradrenergic locus coeruleus (LC) in the brainstem (Figure 1; Supplemental Figure 2). These commonly engaged subcortical areas may constitute an initial circuit recruited by monoaminergic antidepressants, but a single treatment is insufficient to produce a therapeutic effect in patients.

Since a single dose of a traditional monoaminergic antidepressant is subtherapeutic, the above pattern likely reflects early circuit engagement that is inadequate for full antidepressant efficacy. To investigate whether this activation pattern changes with repeated dosing, we next examined the effects of a chronic fluoxetine regimen Mice were treated with fluoxetine (15 mg/kg) daily for two weeks—an early time point at which clinical efficacy of SSRIs typically begins to emerge (Taylor et al., 2006).

After two weeks of daily fluoxetine, we observed enhanced c-fos-based activity across the same subcortical areas identified after a single dose, including the PVT, MD, CEA and BSTa and medial NAc-sh (Figure 1; Supplemental Table 1). Additionally, frontal cortical activation emerged: chronic fluoxetine induced significant c-fos-based activity in the PL and ILA cortex, the dorsal and ventral anterior cingulate cortex (ACCd and ACCv) and the claustrum (CLA), as well as new thalamic activation in the centromedial (CM), reuniens (RE), and rhomboid (RH) nuclei (Figure 1). These findings suggest that recruitment of frontal cortical areas—widely implicated in depression recovery—develops gradually over chronic SSRI treatment. Notably, the MD thalamus has strong reciprocal connectivity with the PL and ILA cortices (Barson et al., 2020; Parnaudeau et al., 2018), raising the possibility that early MD activation (observed after a single dose) contributes to the later engagement of PL/ILA with repeated dosing. Likewise, the medial NAc-sh and the CM, RE, and RH thalamic nuclei all receive prominent inputs from the frontal cortex, suggesting that their robust activation after two weeks of fluoxetine may lie downstream of sustained PL/ILA activity. Taken together, the interconnected network recruited by chronic fluoxetine—comprising five cortical areas (PL, ILA, ACCd, ACCv, CLA), five thalamic nuclei (PVT, MD, CM, RE, RH), and three limbic structures (BSTa, CEA, medial NAc-sh)—may represent a core circuit underlying the antidepressant efficacy of traditional monoaminergic drugs after chronic treatment.

### Ketamine engages the antidepressant circuit acutely and dose-dependently

Because ketamine is effective even in treatment-resistant depression (TRD)—particularly in patients who have not responded to traditional antidepressants—its brain-activity profile might recruit additional regions or stronger activation of the core circuit. To investigate ketamine’s antidepressant effect, we first compared subanesthetic doses of 10 and 30 mg/kg, which have been used in mice to model both its antidepressant action and dissociative side effects (Autry et al., 2011; Schmack et al., 2021; Vesuna et al., 2020).

Whole-brain voxel-wise statistics revealed that both ketamine doses induced c-fos-based activity across a similar pattern of subcortical and cortical areas activated by two-week fluoxetine, with dose-dependent increases most pronounced in the frontal cortical regions and the downstream RE, RH and CM thalamic nuclei and the NAc-sh (Figure 2; Supplemental Table 1). Additionally, ketamine at 10 mg/kg evoked moderate (>200%) and at 30 mg/kg high (>1000%) increases in c-fos-based activity in the dorsal retrosplenial (RSPd) cortex—a region implicated in dissociative perceptions by both mouse and human studies (Parvizi et al., 2021; Tian et al., 2023; Vesuna et al., 2020). The absence of substantial RSPd activation at 10 mg/kg and its dramatic activation at 30 mg/kg, together with previous imaging of ketamine-evoked RSPd activity (Vesuna et al., 2020), suggest that 10 mg/kg represents a low or sub-dissociative dose in mice, whereas 30 mg/kg induces dissociative-like brain activity. Given that ketamine’s rapid antidepressant effects often co-occur with dissociative experiences, the 30 mg/kg dose in mice likely provides a more complete representation of ketamine’s full antidepressant mechanism of action.

**Figure 2.**
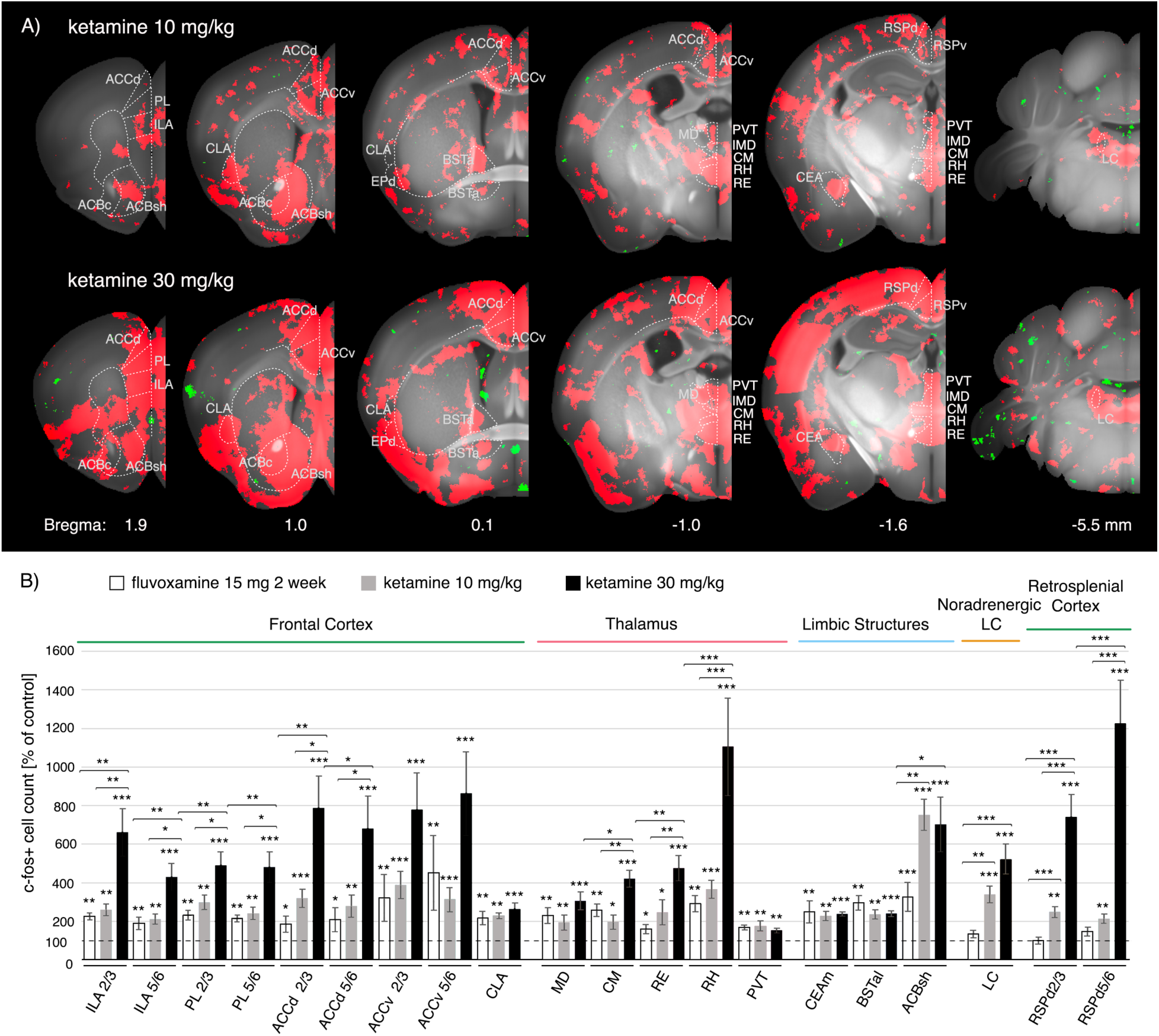
Brain activity evoked by single treatments with subanesthetic ketamine. (**A**) Brain regions with significant c-fos increases (red) are shown on representative coronal sections of the reference mouse brain. *Top:* Pattern evoked by **ketamine** at 10 mg/kg (subanesthetic, sub-dissociative dose). *Bottom:* Pattern evoked by ketamine at 30 mg/kg (higher subanesthetic dose). (**B**) Quantification of c-fos+ cell count changes (mean ± SEM) in key cortical and subcortical structures associated with antidepressant efficacy, for the conditions shown in (A) (10 mg/kg ketamine and 30 mg/kg ketamine) and for a chronic two-week **fluoxetine** treatment (15 mg/kg, FLX 2 wk). (**C**) Quantification of c-fos activation in the **RSPd**—a region linked to ketamine’s dissociative side effects—for ketamine 10 mg/kg, ketamine 30 mg/kg, and chronic fluoxetine (15 mg/kg). (**D**) Direct comparison of chronic fluoxetine (15 mg/kg) and single-dose ketamine (30 mg/kg) treatments on c-fos+ cell counts in selected regions. In (B–D), data are expressed as percent change vs. vehicle controls. Significance in (A–C) was determined by negative binomial regression with FDR correction (q* < 0.05, q** < 0.01, q*** < 0.001). In (D), statistical differences between treatments are indicated by horizontal brackets (one-way ANOVA overall F = 9.22, p < 0.001; post hoc *t*-tests with p* < 0.05, p** < 0.01, p*** < 0.001). See Methods for region abbreviations.

**Figure 3.**
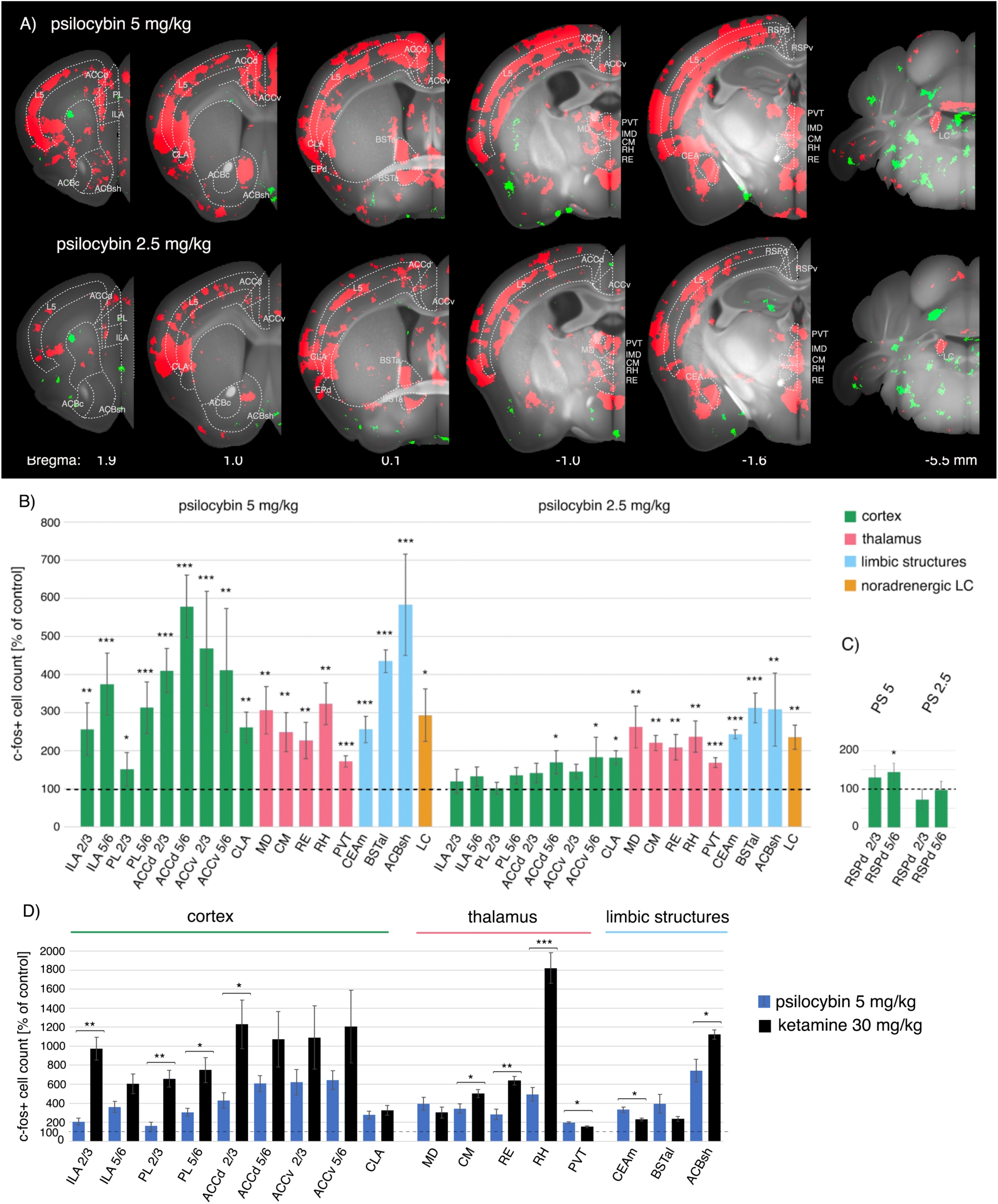
Brain activity evoked by a single treatment with psilocybin. (**A**) Brain regions with significant c-fos increases (red) are shown on coronal sections of the reference brain. *Top:* Pattern evoked by **psilocybin** at 2.5 mg/kg (a subtherapeutic, non-hallucinogenic dose). *Bottom:* Pattern evoked by psilocybin at 5 mg/kg (a therapeutically effective, hallucinogenic dose). (**B**) Quantification of c-fos+ cell changes in key antidepressant-linked regions for the two psilocybin doses (2.5 mg/kg and 5 mg/kg). (**C**) Quantification of c-fos changes in the **RSPd** for psilocybin at 2.5 mg/kg vs. 5 mg/kg, highlighting the lack of RSPd activation at either dose (consistent with psilocybin’s non-dissociative profile). (**D**) Direct comparison of psilocybin (5 mg/kg) and **ketamine** (30 mg/kg) treatments, showing differences in c-fos activation across selected regions (horizontal brackets). Data are expressed as percent change vs. vehicle. Significance in (A–C) was determined by negative binomial regression with FDR correction (q* < 0.05, q** < 0.01, q*** < 0.001). In (D), statistics were determined by one-way ANOVA (overall F = 8.67, p < 0.001) followed by pairwise *t*-tests (p* < 0.05, p** < 0.01, p*** < 0.001). See Methods for abbreviations.

**Figure 4.**
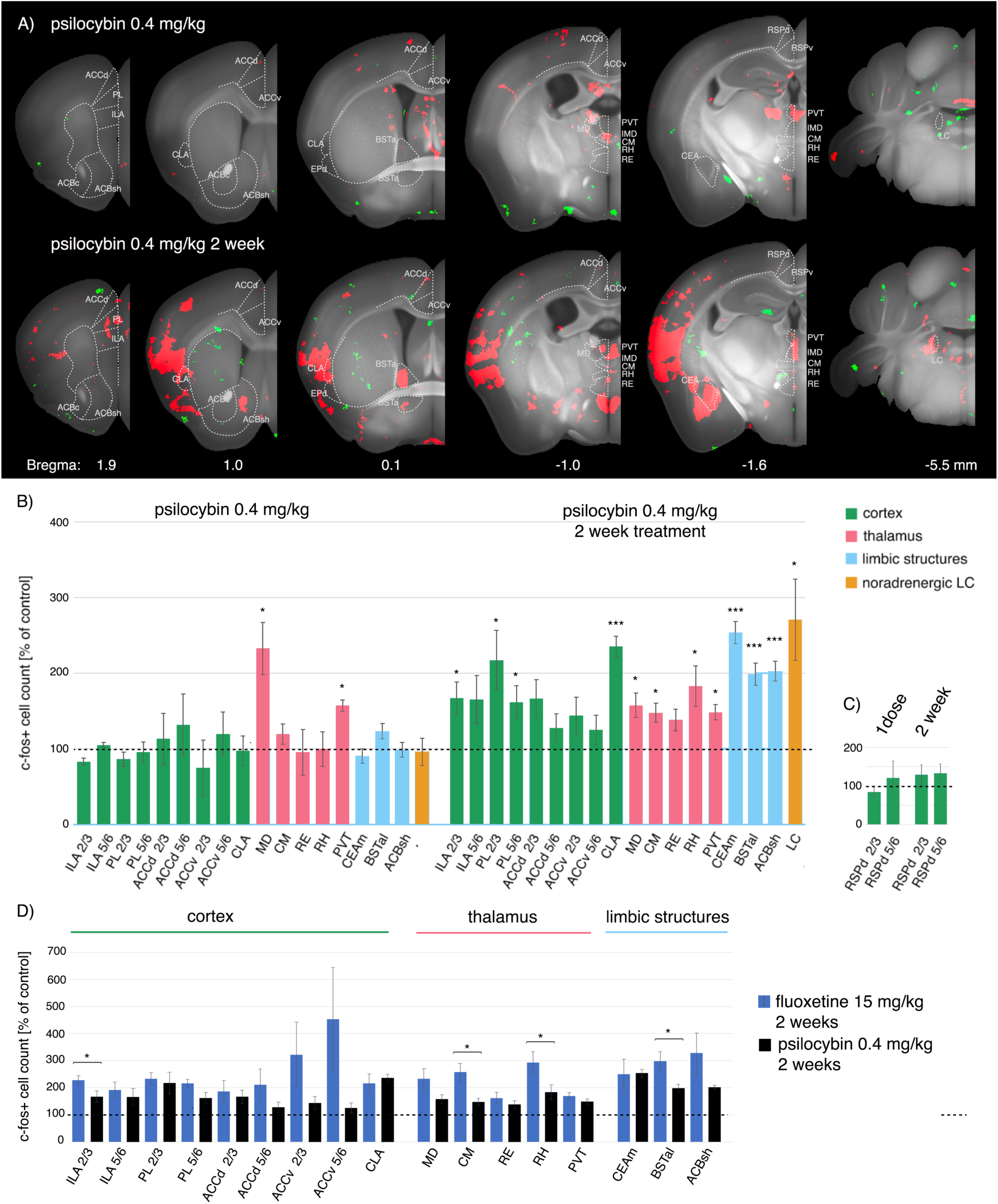
Brain activity evoked by single and chronic treatment with a microdose of psilocybin. (**A**) Brain regions with significant c-fos increases (red) are shown on coronal sections. *Top:* Pattern evoked by a **single psilocybin microdose** (0.4 mg/kg, corresponding to ∼2 mg in humans). *Bottom:* Pattern evoked by a **chronic 2-week psilocybin microdose** regimen (0.4 mg/kg daily). (**B**) Quantification of c-fos+ cell changes in key regions for the single vs. chronic microdose conditions. (**C**) Quantification of c-fos changes in the **RSPd** for single vs. chronic microdose treatment. (**D**) Direct comparison of chronic **fluoxetine** (15 mg/kg, 2 weeks) and chronic psilocybin microdose (0.4 mg/kg, 2 weeks) treatments, highlighting their similar activation patterns with some differences in magnitude (horizontal brackets indicate significant differences). Data are percent change vs. vehicle. Significance in (A–C) determined by negative binomial regression with FDR correction (q* < 0.05, q** < 0.01, q*** < 0.001). In (D), statistics by one-way ANOVA (F = 8.67, p < 0.001) and *t*-tests (p* < 0.05, p** < 0.01, p*** < 0.001). See Methods for abbreviations.

To compare the magnitude of c-fos activation across treatments in specific brain areas, we next quantified c-fos+ cell count changes per anatomical region for 10 and 30 mg/kg ketamine versus two-week fluoxetine (Figure 2). This revealed that 10 mg/kg ketamine and chronic fluoxetine evoked comparable activation in most structures of the putative antidepressant circuit, except in the medial NAc-sh, noradrenergic LC and the RSPd cortex, where 10 mg/kg ketamine evoked significantly c-fos-based higher activity. In contrast, dissociative 30 mg/kg ketamine evoked markedly greater (2-3-fold) activity than two-week fluoxetine or 10 mg/kg ketamine in the frontal cortical PL, ILA and ACCd cortex, the RSPd cortex, the thalamic CM, RE and RH nuclei and the medial NAc-sh. Notably, no significant differences in c-fos-based activity were observed among the three treatments (fluoxetine vs. 10 mg vs. 30 mg ketamine) in the MD and PVT thalamus or in the CEA and BSTa—all regions that were activated even by a single SSRI/TCA dose.

To further examine ketamine’s dose–response characteristics, we tested an even lower dose and a much higher dose. A low ketamine dose (5 mg/kg) failed to induce any significant c-fos-based activity aside from a distinct hotspot in the medial NAc-sh, suggesting that this region is among the most sensitive to ketamine’s effects. In contrast, an anesthetic-level dose (120 mg/kg) evoked widespread cortical activation encompassing nearly all cortical areas, far beyond the more selective cortical pattern seen with subanesthetic doses (Supplemental Figure 3).

Comparing across doses (5, 10, 30, 120 mg/kg) revealed that the highest dose produced maximal c-fos-based activity primarily in the f frontal cortical areas, RSPd, and the midline thalamic nuclei CM, RE, RH, with only moderate or no further increase in the MD, PVT, CEA, or BSTa beyond the subanesthetic doses and two-week fluoxetine.

Taken together, these experiments reveal structure-specific distinctions in ketamine’s activation profile: while most cortical regions and several thalamic nuclei exhibited the expected dose-dependent increases in c-fos-based activity, the thalamic PVT and MD and the limbic CEA and BSTa—regions already activated by single-dose monoaminergic antidepressants—showed flat dose–response curves (no additional recruitment at higher ketamine doses).

### Psilocybin engages the antidepressant circuit at a high dose

Psilocybin is being actively investigated in clinical trials for depression. Early studies indicate that a single hallucinogenic dose (25 mg in humans) can produce rapid antidepressant effects lasting weeks, whereas a non-hallucinogenic dose (10 mg) does not confer clear therapeutic benefit (Davis et al., 2021; Goodwin et al., 2022; Perez et al., 2023; Raison et al., 2023). We sought to determine whether psilocybin engages the same core brain structures as ketamine or chronic fluoxetine, or whether it recruits distinct circuits. Mice were treated with psilocybin at 2.5 mg/kg or 5 mg/kg, doses approximating the human 10 mg and 25 mg doses, respectively.

The higher, hallucinogenic dose (5 mg/kg) of psilocybin evoked robust activation across all subcortical brain structures activated by ketamine and chronic fluoxetine, as well as widespread cortical activity that included the PL, ILA, and ACC but also all sensory cortical areas (Figure 3). This cortical activity was especially pronounced in layer 5, which comprises pyramidal neurons with high expression of 5HT2A receptors believed to mediate psilocybin’s hallucinogenic effects (Pompeiano et al., 1994). Psilocybin at 5 mg/kg, however, failed to evoked significant activation of the RSPd cortex (Figure 3), consistent with its lack of dissociative effects. A direct comparison of psilocybin 5 mg/kg and ketamine 30 mg/kg revealed clear differences. Ketamine induced significantly higher c-fos-based activity than psilocybin in the PL and ILA cortices, as well as in the CM, RE, and RH thalamic nuclei and the medial NAc-sh. Conversely, psilocybin induced significantly higher c-fos activity than ketamine in the CEA and PVT (Figure 3). These distinct patterns suggest that while both drugs converge on a common limbic–frontal circuit, ketamine drives stronger activity in the frontal cortex, whereas psilocybin evokes greater activation in subcortical limbic regions, along with widespread layer 5 cortical activity.

The lower psilocybin dose (2.5 mg/kg) produced a notably different pattern. It failed to evoke significant c-fos induction in the frontal PL and ILA areas, yet still produced robust activation of all the subcortical limbic structures, including the BSTa, CEA, NAc-sh, thalamic nuclei, and the LC (Figure 3).

The lower dose of 2.5 mg/kg psilocybin failed to evoke significant activity in the frontal cortical PL and ILA areas, but retain robust activation of all subcortical structures, including BSTa, CEA, NAc-sh, the midline thalamic nuclei, and LC (Figure 3). Since at a subtherapeutic dose, psilocybin activated the same subcortical circuit as the monoaminergic drugs, but without engaging the frontal cortical nodes of the circuit, this suggests that recruitment of the PL/ILA cortex may be a necessary step for rapid-acting antidepressant efficacy.

Psilocybin’s active metabolite, psilocin, has the highest affinity at the 5HT2A receptors but also acts at 5HT2C and 5HT1A receptors (Erkizia-Santamaría et al., 2022). To investigate the contributions of 5-HT2A/2C versus 5-HT1A receptors to psilocybin’s c-fos-based activity, we examined the activity of the hallucinogen 2C-T, which acts primarily at 5-HT2A and 5-HT2C receptors (Luethi et al., 2018). Administration of 2C-T at 35 mg/kg (approximating a hallucinogenic human dose of ∼170 mg (Shulgin and Shulgin, 1991)) evoked a layer 5-centric cortical activation similar to that of psilocybin, including strong c-fos induction in the PL, ILA, and ACC (Supplemental Figure 4A). 2C-T also activated the limbic subcortical structures BSTa, CEA, and NAc-sh, but notably failed to activate several midline thalamic nuclei (including the MD, PVT, and RE) that were significantly activated by ketamine, chronic fluoxetine, and hallucinogenic-dose psilocybin (Supplemental Figure 4). These findings suggest that 5-HT2A receptor activation is primarily responsible for the distinctive layer 5 cortical activity pattern, whereas engagement of both 5-HT2A and 5-HT1A receptors may be needed to produce the combined cortical-plus-subcortical activation seen with psilocybin.

### Psychedelic microdosing converges on the antidepressant circuit with repeated use

Microdosing—the practice of taking very low, non-hallucinogenic doses of psychedelics—has been anecdotally reported to improve mood, perception, and cognition. Typically, a psilocybin “microdose” is about 1–3 mg (e.g., consuming 0.2–0.5 g of dried Psilocybe mushrooms).

However, placebo-controlled studies in healthy volunteers have so far failed to demonstrate clear, clinically relevant benefits of microdosing, and no data from controlled trials in depression patients are yet available (Cavanna et al., 2022; Rootman et al., 2022). We investigated psilocybin microdosing in our mouse assay by comparing c-fos-based activaty after a single microdose versus chronic microdosing. Mice received psilocybin at 0.4 mg/kg (∼2 mg human equivalent) either as a one-time treatment or daily for two weeks.

A single 0.4 mg/kg psilocybin microdose induced significant c-fos-based activity only in the thalamic PVT and MD nuclei (Figure 4). By contrast, two weeks of daily microdosing resulted in a potentiated activation of the subcortical circuit along with moderate activation emerging in frontal cortical regions and the medial NAc-sh (Figure 4). In fact, the pattern of c-fos-based activity after two-week microdosing was broadly similar to that seen with two-week fluoxetine: a direct comparison revealed that chronic psilocybin microdosing produced an overall similar pattern of c-fos-based activation to chronic fluoxetine (Figure 4), albeit at modestly lower levels in the ILA cortex, CM and RH thalamic nuclei, and BSTa. These results demonstrate that psilocybin microdosing, when administered daily, can gradually recruit the same antidepressant-linked circuit as chronic SSRI treatment. This convergence is perhaps unsurprising given that both psilocybin and SSRIs act to enhance serotonergic signaling in the brain. Notably, our study employed a daily dosing schedule for direct comparison with fluoxetine, whereas real-world microdosing regimens are often intermittent (e.g., one day on, one day off, or one day on, two days off). Such intermittent dosing has been hypothesized to optimize efficacy while minimizing tolerance, so it remains to be determined whether non-daily microdosing might produce stronger effects on this circuit.

## Discussion

In this study, we employed unbiased whole-brain c-fos mapping in mice to identify a shared set of interconnected subcortical and frontal cortical regions activated by a two-week fluoxetine or psilocybin microdosing treatment, as well as single treatments with subanesthetic ketamine or hallucinogenic psilocybin. Since chronic SSRI treatment and acute ketamine provide therapeutic relief for depression, these brain structures represent candidate nodes of a core antidepressant-responsive circuit.

Notably, each treatment engages distinct molecular targets broadly distributed across the brain: the serotonin presynaptic transporter (SERT) for fluoxetine, the NMDA receptor for ketamine, and the 5HT2A, 5HT2C, and 5HT1A serotonin receptors for psilocybin. The observed increases in c-fos+ cell counts within a specific set of brain areas suggests a downstream circuit-level convergence of these initially disparate pharmacological actions, resulting in a lasting neural activity and plasticity (as marked by c-fos expression and new protein synthesis) in a common network of brain regions.

The identified regions form a closely interconnected network that can be conceptually subdivided into two subcircuits based on their activation profiles. One subcircuit—comprising the PVT and MD thalamic nuclei along with the BSTa and CEA in the extended amygdala—is activated even by single, acute doses of all the tested traditional monoaminergic antidepressants (Figure 5). The second subcircuit includes the PL and ILA cortices (rodent homologues of the human pgACC and sgACC) as well as the medial NAc-sh, all of which have been implicated in the pathophysiology of depression through human imaging studies (Alexander et al., 2021; Drevets et al., 2008; Lynch et al., 2024). The frontal cortical subcircuit was not engaged by an acute SSRI or low-dose psilocybin, but it was gradually recruited over the course of two-week SSRI or psilocybin microdosing treatment (Figure 5A).

**Figure 5.**
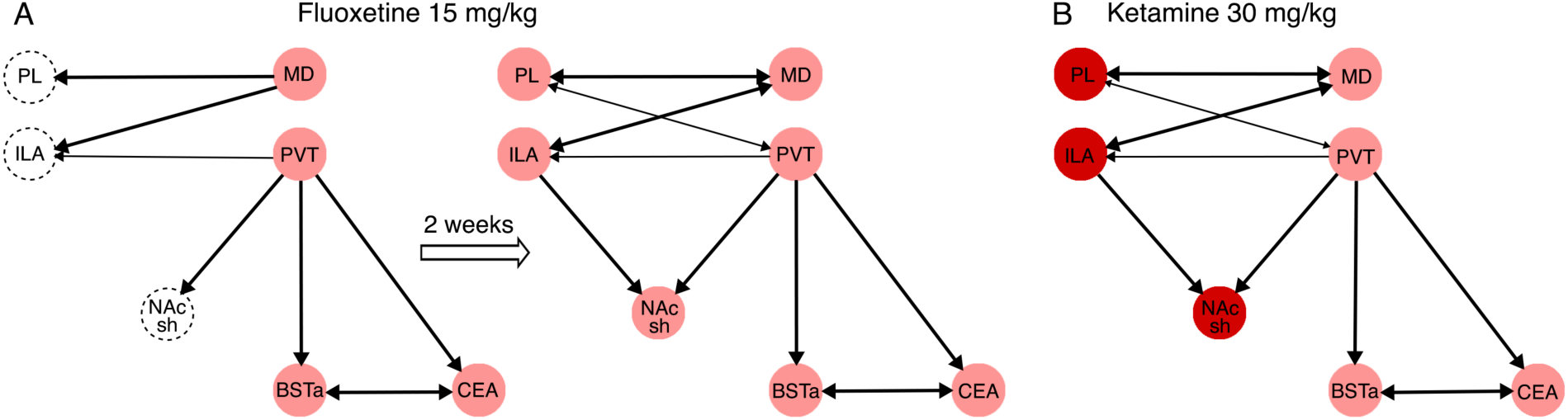
Schematic of an antidepressant-linked brain circuit. This schematic summarizes the core limbic–cortical circuitry that is engaged by effective antidepressant treatments (see Discussion for details). (**A**) Left: A single acute **fluoxetine** (15 mg/kg) treatment activates an initial subcortical circuit involving the thalamic MD and PVT nuclei and the limbic BSTa and CEA. Right: Following a chronic 2-week fluoxetine treatment, the circuit expands to include significant activation of the frontal cortical PL and ILA areas and the medial NAc-sh. (**B**) A single subanesthetic **ketamine** treatment (30 mg/kg) evoked c-fos-based activity across all the structures engaged by the chronic fluoxetine treatment. Arrows indicate major efferent projections between the activated regions; thicker arrows denote particularly strong or dense connections.

The MD thalamus plays a key role in emotion, motivation, and executive function. It integrates inputs from the basolateral amygdala and limbic forebrain and serves as a principal thalamic relay to the PL, ILA and other frontal cortical areas (Parnaudeau et al., 2018). The PVT primarily integrates arousal- and stress-related signals from the brainstem and relays them to the extended amygdala stress circuitry, including the reciprocally connected BSTa and CEA (Barson et al., 2020). Additionally, PVT contributes to reward and motivation processing through its projections to the NAc-sh and, to a lesser extent, the PL and ILA cortices—although its cortical projections are sparser than those of the MD thalamus (Kark et al., 2021). The consistent activation of the PVT-MD-BSTa-CEA subcircuit by single doses of antidepressants, combined with its connectivity to the PL and ILA, suggests it may play an essential role in initiating the antidepressant response. We speculate that early activation of this thalamo-amygdala pathway could gradually drive the recruitment of frontal cortical regions, ultimately leading to the sustained frontal cortical activation associated with therapeutic antidepressant effects.

Subanesthetic ketamine, while rapidly effective, frequently induces dissociative experiences in patients, necessitating administration under medical supervision. Imaging and electrophysiological studies in both mice and humans have identified a characteristic 1–3 Hz rhythm in the RSPd and its human homologue the posteromedial cortex as a signature of ketamine’s dissociative effects (Tian et al., 2023; Vesuna et al., 2020). This 1–3 Hz oscillation in the RSPd is observed at a 30 mg/kg ketamine dose but not at 10 mg/kg in mice (Vesuna et al., 2020). Consistent with that, our study found a striking (>1000%) increase in c-fos-based activity in the RSPd at 30 mg/kg but only moderate (200%) increased at 10 mg/kg. Taken together, these data indicate that 10 mg/kg is a sub-dissociative dose in mice, while 30 mg/kg more accurately reflects ketamine’s full antidepressant efficacy in TRD.

Comparing ketamine (10 vs. 30 mg/kg) with chronic fluoxetine provided further insight. Overall, we found that ketamine at 10 mg/kg activated the putative antidepressant circuit to a similar degree as two-week fluoxetine. Surprisingly, raising the ketamine dose to 30 mg/kg did not increase c-fos activity in the MD, PVT, BSTa, or CEA—suggesting that these regions may already be maximally engaged at the lower dose. In contrast, the PL and ILA cortices and the RH, RE, CM thalamic nuclei showed 2–3-fold higher activity at 30 mg/kg than at 10 mg/kg.

These findings suggest a model in which the MD–PVT–BSTa–CEA subcircuit acts as a common gateway for inducing an antidepressant response (being activated by even low doses and conventional drugs), whereas the strength or completeness of the antidepressant effect is reflected in the degree of frontocortical and downstream midline thalamic activation. In the case of ketamine, the much greater recruitment of PL/ILA and midline thalamus at 30 mg/kg may be linked to its enhanced efficacy in TRD. Indeed, the increased c-fos-based activity we observed between 10 and 30 mg/kg mirrors dose-dependent increases in clinical antidepressant efficacy reported for ketamine between 0.1 and 0.5 mg/kg i.v. in humans (Seshadri et al., 2024).

At an anesthetic dose (120 mg/kg), ketamine induced an essentially global wave of cortical c-fos expression, along with further potentiation of the frontocortical, RSPd, and RH, RE, CM thalamic activity. This broad activation likely corresponds to the state of dissociative anesthesia, in which a patient appears awake (eyes open) but is unresponsive to pain or external stimuli. Interestingly, clinical studies have found that anesthetic-dose ketamine (≥2 mg/kg i.v.) fails to enhance the efficacy of electroconvulsive therapy (ECT) or to alleviate depressive symptoms when administered during major surgery ((Mashour et al., 2018; Rasmussen et al., 2014). This suggests that the more spatially selective pattern of activation seen with subanesthetic ketamine—focusing on a specific frontal–limbic circuit rather than the entire cortex—is critical for ketamine’s antidepressant effects.

The medial NAc-sh, in particular the rostral–dorsal subregion, integrates motivational valence and processes novel or salient stimuli, and its dysregulation has been linked to the symptoms of anhedonia and avolition in depression (Pizzagalli et al., 2009). In our study, a single dose of fluoxetine did not significantly activate the medial NAc-sh, whereas two-week fluoxetine produced a robust, localized hotspot of c-fos-based activity in this region. Given that the NAc-sh receives converging excitatory inputs from both the PL/ILA cortex and the PVT, this delayed activation may reflect the integration of top-down frontal signals with thalamic arousal signals following chronic SSRI exposure. In contrast, even a very low ketamine dose (5 mg/kg), which produced minimal activation elsewhere, elicited a similar hotspot in the medial NAc-sh.

This might indicate that a subthreshold engagement of PL/ILA and PVT—insufficient to drive widespread c-fos on their own—can nonetheless synergize at their convergence point in the NAc-sh to produce a detectable signal. Notably, the subregion of NAc-sh we identified overlaps with a known “hedonic hotspot” that, when stimulated, amplifies the pleasure (“liking”) response to rewarding stimuli such as sucrose (Castro and Berridge, 2014),. This finding supports a mechanistic link between activation of this PL/ILA–PVT–NAc circuit and the alleviation of anhedonia in depression.

Additional brain regions activated by both chronic fluoxetine treatment and subanesthetic ketamine include the ACCd and ACCv, CLA, and the thalamic nuclei RH, RE, and CM. In rodents, ACCd and ACCv are adjacent to PL/ILA as part of the medial prefrontal cortex and are involved in integrating cognitive, emotional, and pain-related information. The claustrum is one of the most highly connected structures in the brain, with reciprocal connections to nearly all cortical areas. Its putative functions include integrating sensory, cognitive, and emotional information to coordinate attention and adaptive behavior. The RE and RH)nuclei provide a bridge between the medial prefrontal cortex (PL/ILA) and the hippocampal formation, forming circuits important for memory, attention, and emotional regulation (Cassel et al., 2021; Vertes et al., 2015). The intralaminar CM)nucleus is primarily connected with the PL cortex and is implicated in arousal and attention. Activation of this broad supporting cast of regions by both SSRI and ketamine suggests they are integral components of the extended antidepressant network, potentially contributing to cognitive-emotional integration and enhanced information processing as depression remits.

Turning to psilocybin, we found that while it activates the same core circuit, its pattern of cortical engagement is clearly distinct from that of ketamine or monoaminergic drugs. A hallucinogenic dose of psilocybin (5 mg/kg in mice, ∼25 mg in humans) induced widespread cortical activation, especially in layer 5 pyramidal neurons that express 5-HT 2A receptors (Davis et al., 2021; Goodwin et al., 2022; Raison et al., 2023). This broad cortical engagement likely underlies psilocybin’s profound alterations of perception and cognition (hallucinogenic effects). In our data, the activation of the PL and ILA cortices by psilocybin was present but considerably weaker than that observed with ketamine. This suggests that the antidepressant efficacy of psilocybin (at hallucinogenic doses) may depend, at least in part, on its diffuse cortical activation in addition to whatever engagement of PL/ILA does occur. In other words, psilocybin’s antidepressant effect might rely on a more distributed cortical plasticity, rather than strongly targeting the mPFC alone—though further studies (including ongoing phase III trials) will be needed to confirm this. Furthermore, the fact that a lower, non-hallucinogenic dose of psilocybin (2.5 mg/kg, ∼10 mg human) produced robust subcortical activation without engaging PL/ILA provides additional support for the hypothesis that frontocortical recruitment is a key determinant of antidepressant action. Even though the subcortical limbic circuit was activated at 2.5 mg/kg, the lack of PL/ILA activation in that condition correlated with its known lack of therapeutic efficacy.

Finally, our results with psilocybin microdosing revealed the following notable. A single microdose in mice (0.4 mg/kg) induced significant activation only in the MD and PVT thalamic nuclei, yet chronic microdosing produced a circuit activation pattern strikingly analogous to that of chronic fluoxetine (albeit slightly attenuated). This convergence implies that psilocybin microdosing may work through the same brain pathways as traditional antidepressants, by gradually amplifying activity in the limbic–frontal circuit. This finding dovetails with the idea that both SSRIs and low-dose psychedelics boost serotonin signaling, ultimately impinging on a shared neural substrate. While these data suggest that microdosing may not offer additional advantage over traditional antidepressant, it is important to note that our protocol differed from many real-world microdosing practices. Intermittent dosing schedules (e.g., dosing every other day) might elicit stronger cumulative effects or reduce tolerance, potentially enhancing the impact on this circuit beyond what we observed with daily dosing. Future studies that vary the microdosing schedule could address this possibility.

In summary, our findings define a set of interconnected frontal cortical and subcortical limbic structures as the core neural substrates of antidepressant efficacy in the mouse brain. This model helps explain the mechanistic basis for the delayed effects of traditional monoaminergic antidepressants versus the rapid action of ketamine, and it suggests that traditional antidepressants, ketamine, and psilocybin ultimately converge on shared neural circuitry to mediate their antidepressant actions. Identifying this common pathway provides a framework for understanding how diverse treatments relieve depression, and it may inform the development of novel interventions that more directly target this core circuit to achieve faster and more robust antidepressant outcomes.

## Acknowledgment/Funding

We thank Adam Kepecs, Brett Mensh, John Krystal, and Jessica Londa for critical comments on the manuscript. Theracast in a biotechnology company with R&D focused on psychiatric drug discovery. This work was supported by Theracast.

---

**Supplemental Figure 1.**
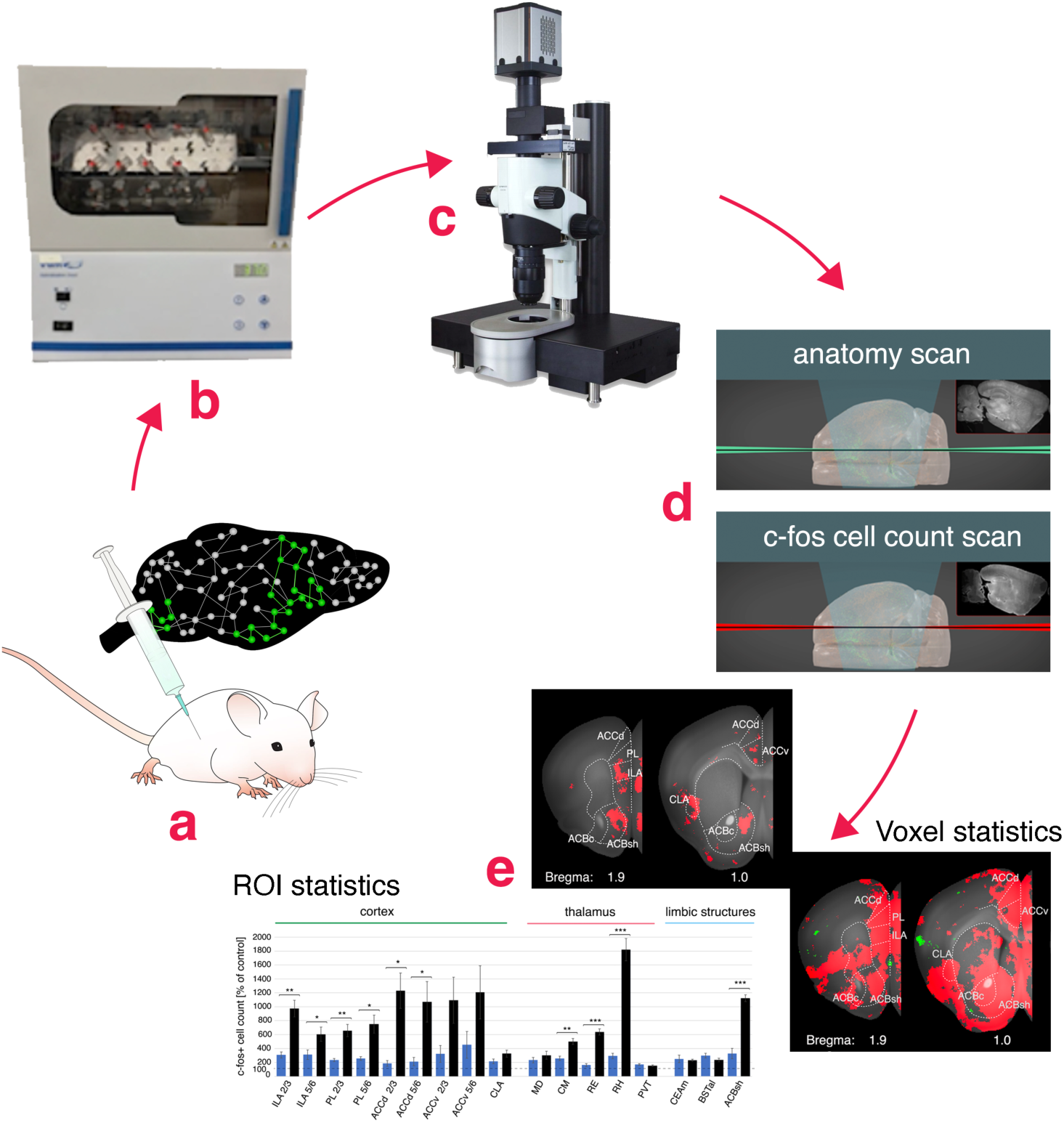
**C-fos mapping pipeline for unbiased drug screening across the mouse brain at single-cell resolution. (**A) Mice receive either a drug or vehicle control via intraperitoneal (i.p.) injection. The drug’s effects on brain cells and circuits may lead to changes in c-fos expression, marking brain areas undergoing lasting drug-induced plasticity (schematically highlighted as a green neural circuit). (B) After a 2.5-hour post-treatment rest in the home cage to allow for c-fos mRNA and protein expression, mice are sacrificed and their brains are extracted and immunolabeled for c-fos protein (via a modified iDISCO+ whole-brain immunolabeling and clearing protocol; see Methods). (C) The cleared brains are imaged with light-sheet fluorescence microscopy (LSFM) to capture both an autofluorescence anatomical scan (488 nm excitation) for anatomical reference and a c-fos immunofluorescence scan (640 nm) to visualize hundreds of thousands of c-fos+ cell nuclei per hemisphere. (D) Each brain’s autofluorescence image is registered to a common 3D reference brain. Computational detection of c-fos+ cells allows mapping of c-fos signals onto the reference, enabling quantitative comparisons across brains. (E) Two complementary statistical analyses are performed: (1) voxel-based analysis, which unbiasedly identifies spatial clusters of significant c-fos+ cell count differences across the entire brain (results visualized in red on the reference brain, as in Figures 1–4), and (2) region-of-interest (ROI) analysis, which quantifies c-fos+ differences as mean ± SEM per annotated anatomical region (as in bar graphs in Figures 1–4). Both analyses use negative binomial regression with FDR correction for multiple comparisons (see Methods for details).

**Supplemental Figure 2.**
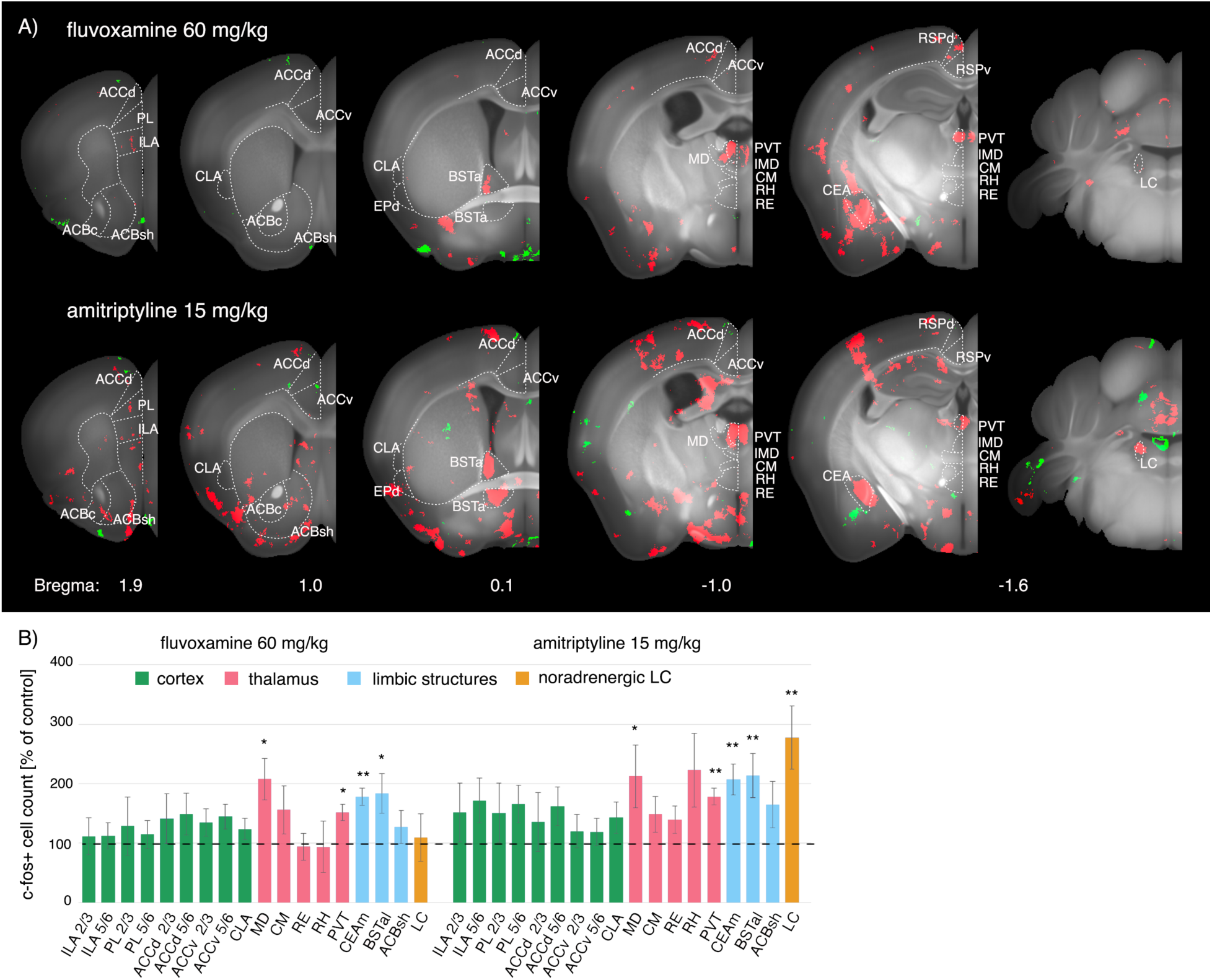
B**r**ain **activity evoked by single treatments with monoaminergic antidepressants.** (**A**) Brain regions with significant c-fos increases (red) are shown on representative coronal sections of the reference brain. *Top:* Pattern evoked by a single dose of the SSRI **fluvoxamine** (60 mg/kg). *Middle:* Pattern evoked by a single dose of the TCA **amitriptyline** (15 mg/kg). *Bottom:* Pattern evoked by a single dose of the atypical antidepressant **mirtazapine** (15 mg/kg). (**B**) Quantification of c-fos+ cell changes (percent vs. vehicle, mean ± SEM) in brain regions linked to antidepressant activity, for the treatments shown in (A). See Supplemental Methods for region abbreviations. (Data for single fluoxetine and desipramine are shown in Figure 1.)

**Supplemental Figure 3.**
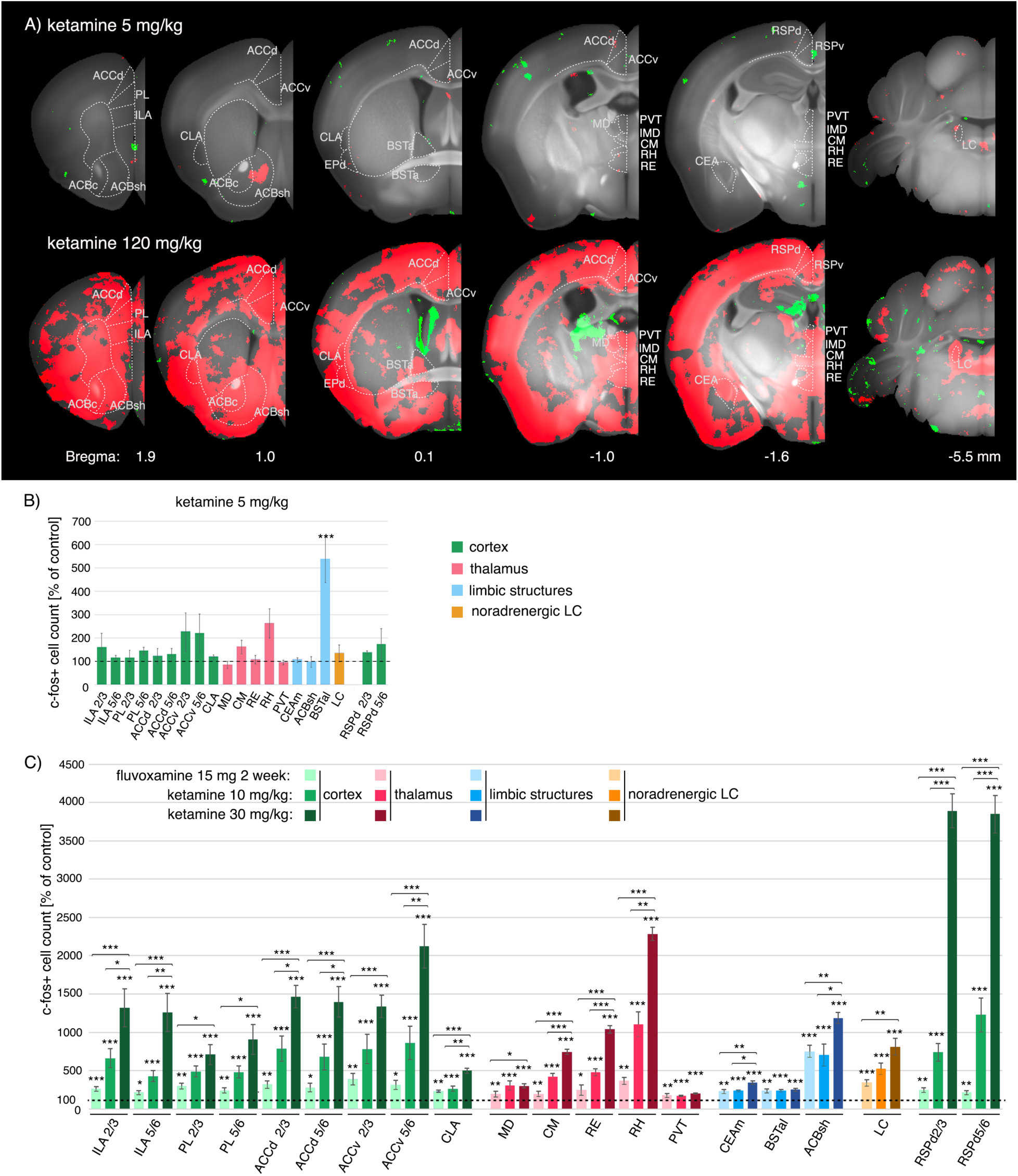
B**r**ain **activity evoked by a single treatment with high-dose (anesthetic) ketamine.** (**A**) Brain regions with significant c-fos increases (red) are shown on coronal sections for **ketamine** at 120 mg/kg (an anesthetic dose). (**B**) Quantification of c-fos+ cell changes (mean ± SEM) in selected cortical and subcortical structures associated with antidepressant efficacy, for ketamine 120 mg/kg. For comparison, data for ketamine 10 mg/kg, ketamine 30 mg/kg, and chronic fluoxetine (15 mg/kg) are also shown (light gray bars; see Figure 2). Data are percent change vs. vehicle. Region abbreviations are defined in Methods.

**Supplemental Figure 4.**
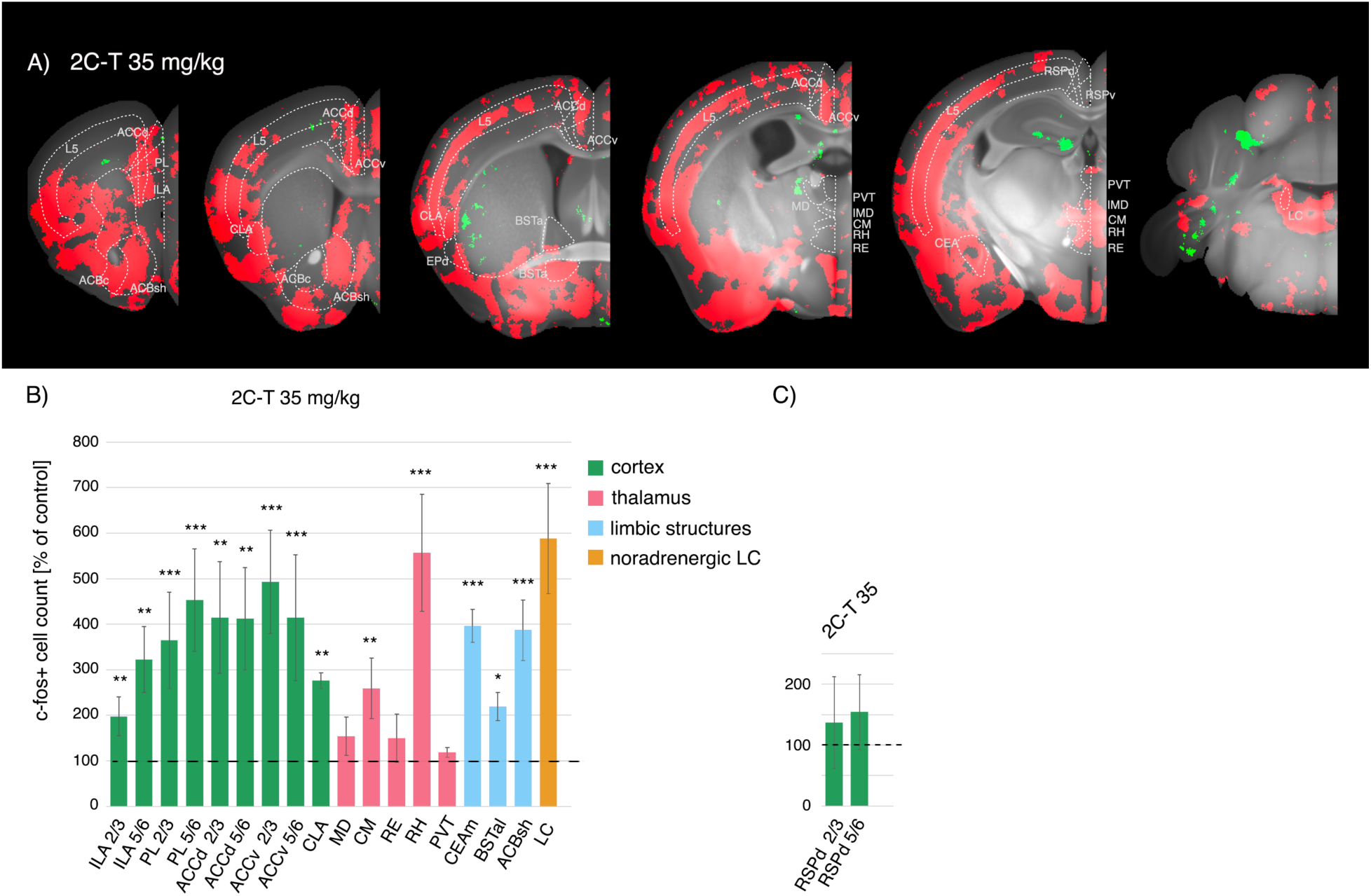
Brain activity evoked by a single treatment with the hallucinogen 2C-T. (A) Brain regions showing statistically significant increases in c-fos expression evoked by a single dose of 2C-T (35 mg/kg) were detected using voxel-based statistics and are visualized in red, overlaid on selected coronal planes from the 3D Reference LSFM mouse brain used for data registration (see Methods). (B) Quantification of the evoked activity shown in (A), across brain regions putatively linked to antidepressant efficacy. (C) Quantification of the evoked activity in the retrosplenial cortex (RSPd) shown in (A), putatively linked to dissociative side effects. The abbreviations used are described in the Supplemental Methods.

